## Supplementary figures and images for "Intracellular Lipid Droplet Accumulation Occurs Early Following Viral Infection and Is Required for an Efficient Interferon Response"

### Supplementary figure 1

#### Supplementary Figure 1

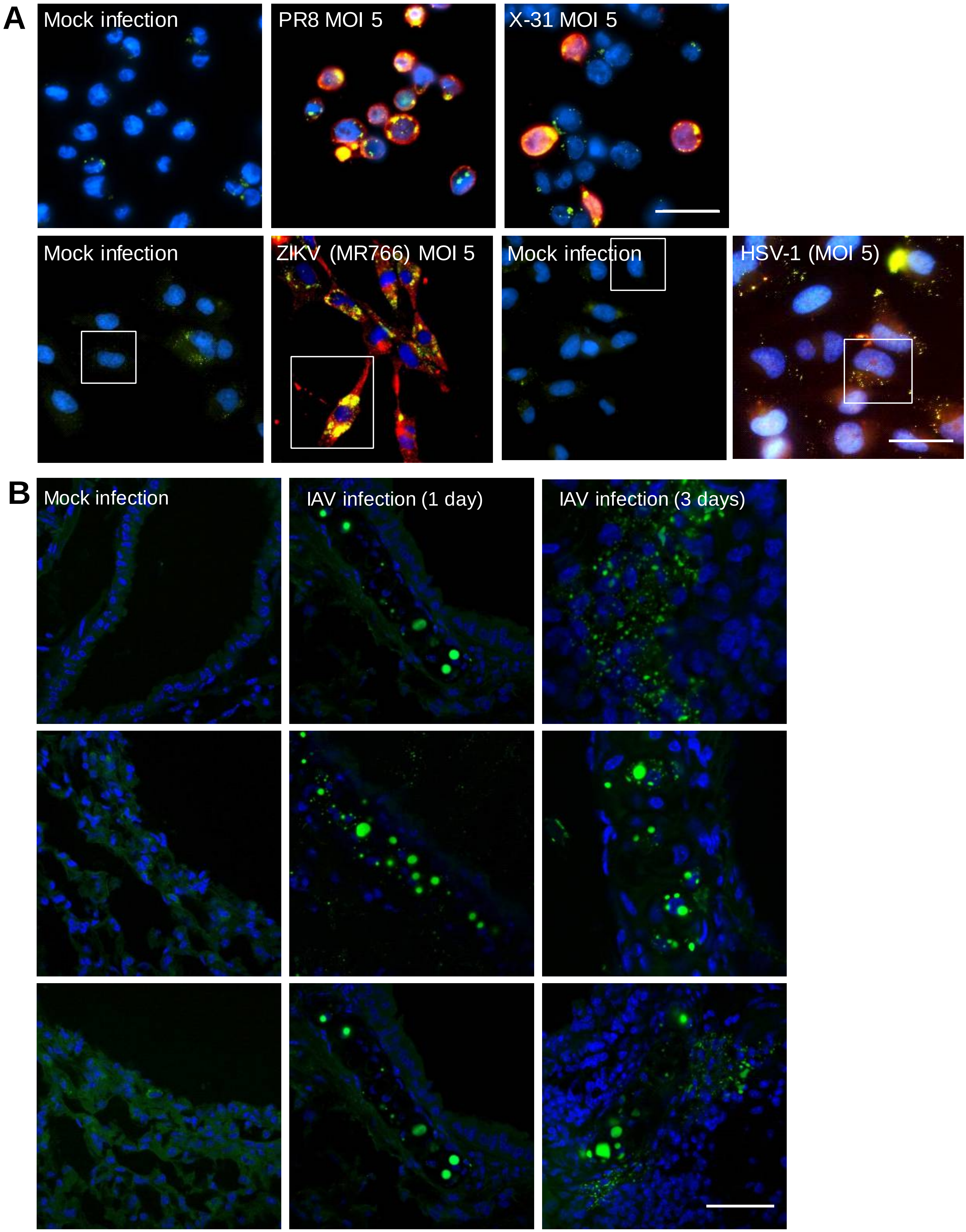

### Supplementary figure 2

#### Supplementary Figure 2

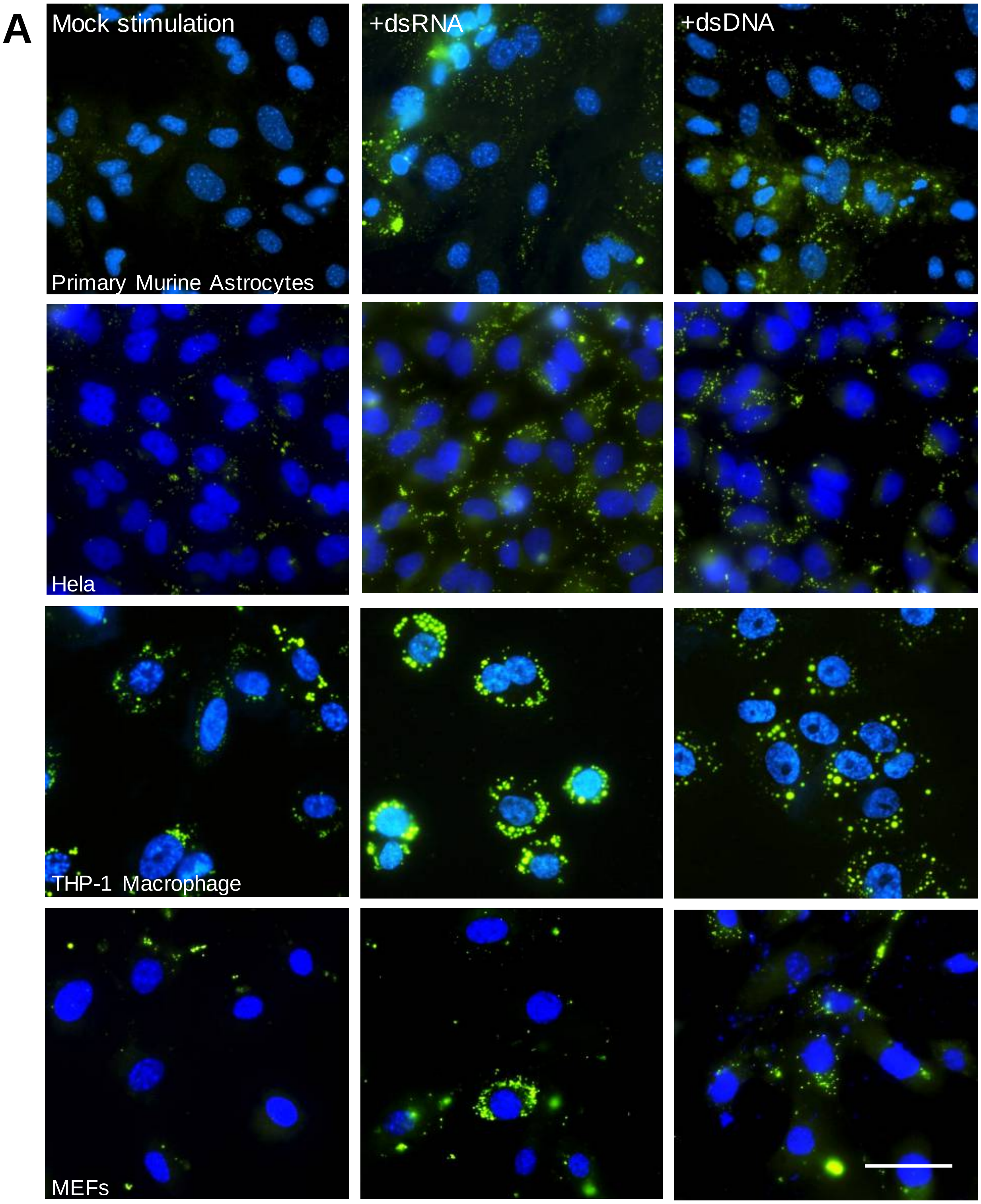


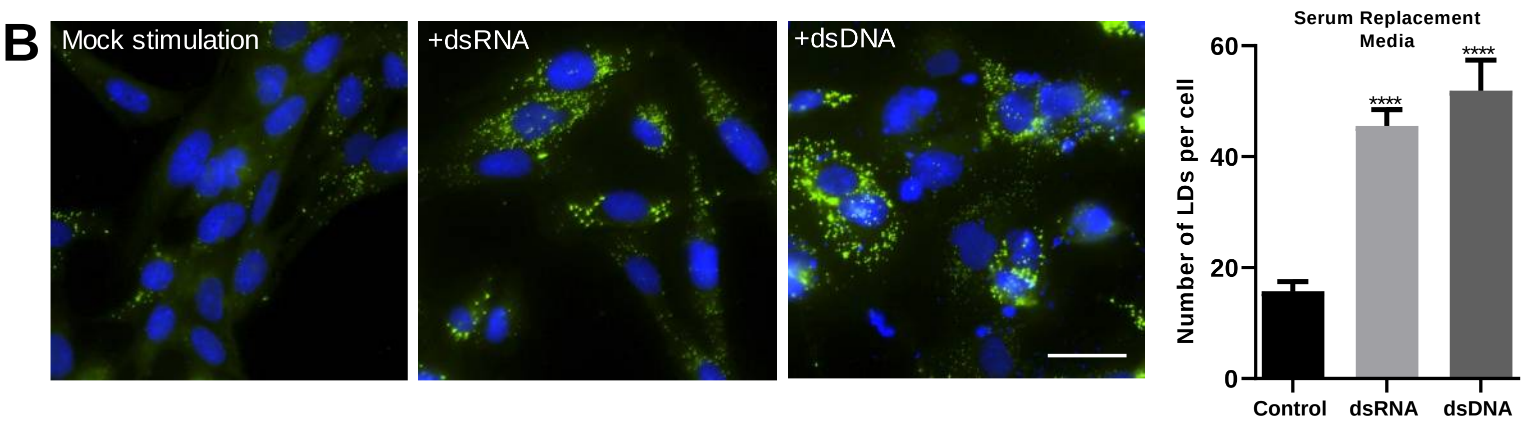

### Supplementary figure 3

#### Supplementary Figure 3

**A**

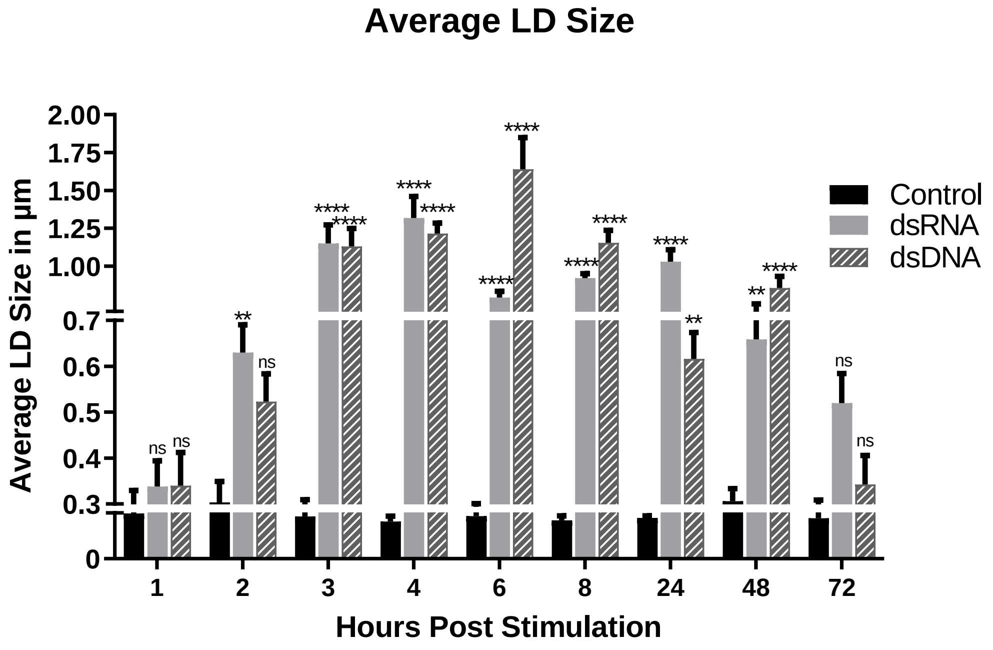

### Supplementary figure 4

#### Supplementary Figure 4

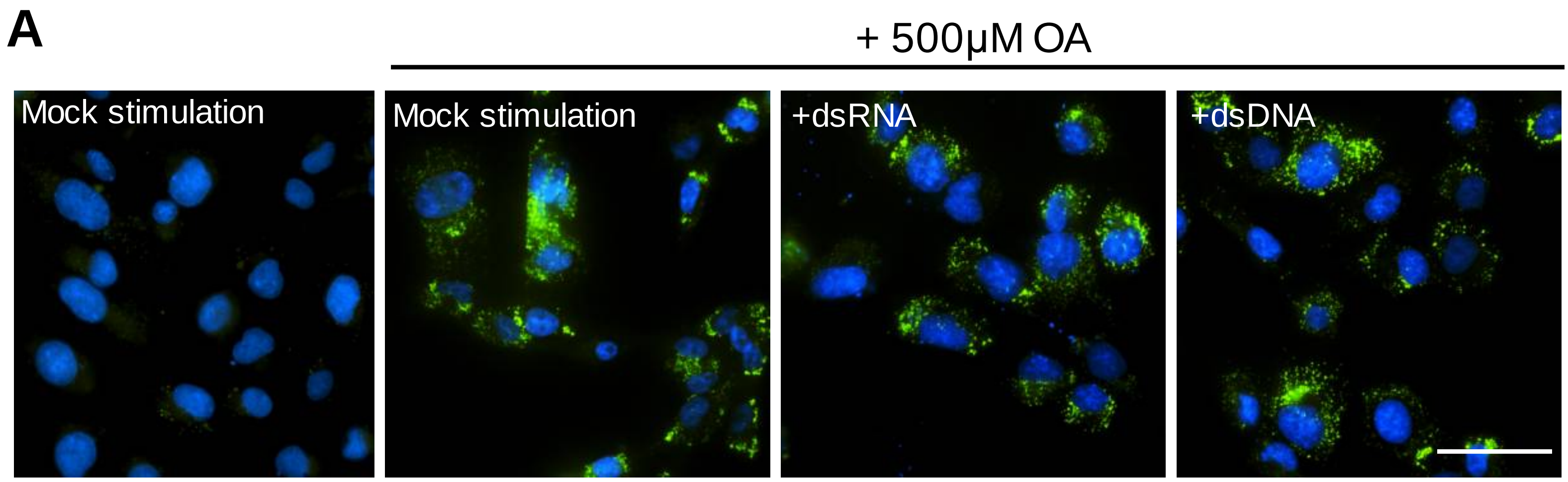

### Supplementary figure 5

#### Supplementary Figure 5

**A**

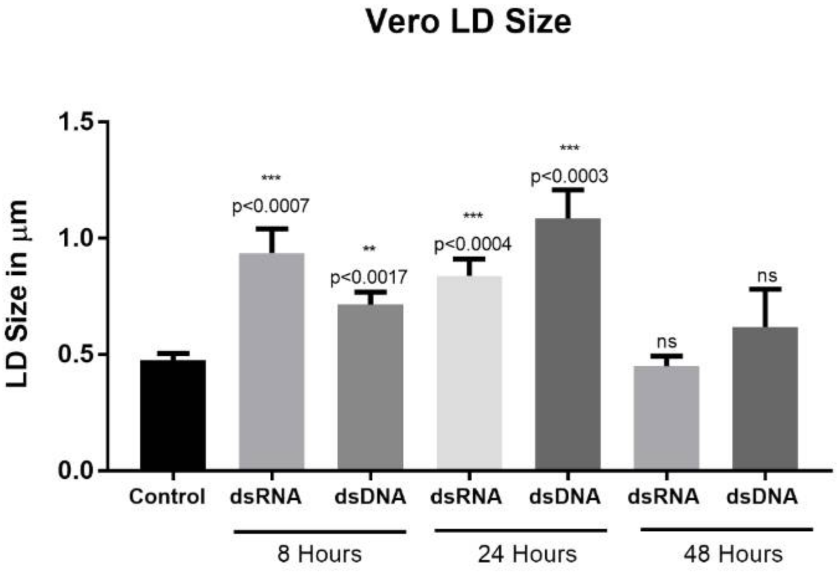

### Supplementary figure 6

#### Supplementary Figure 6

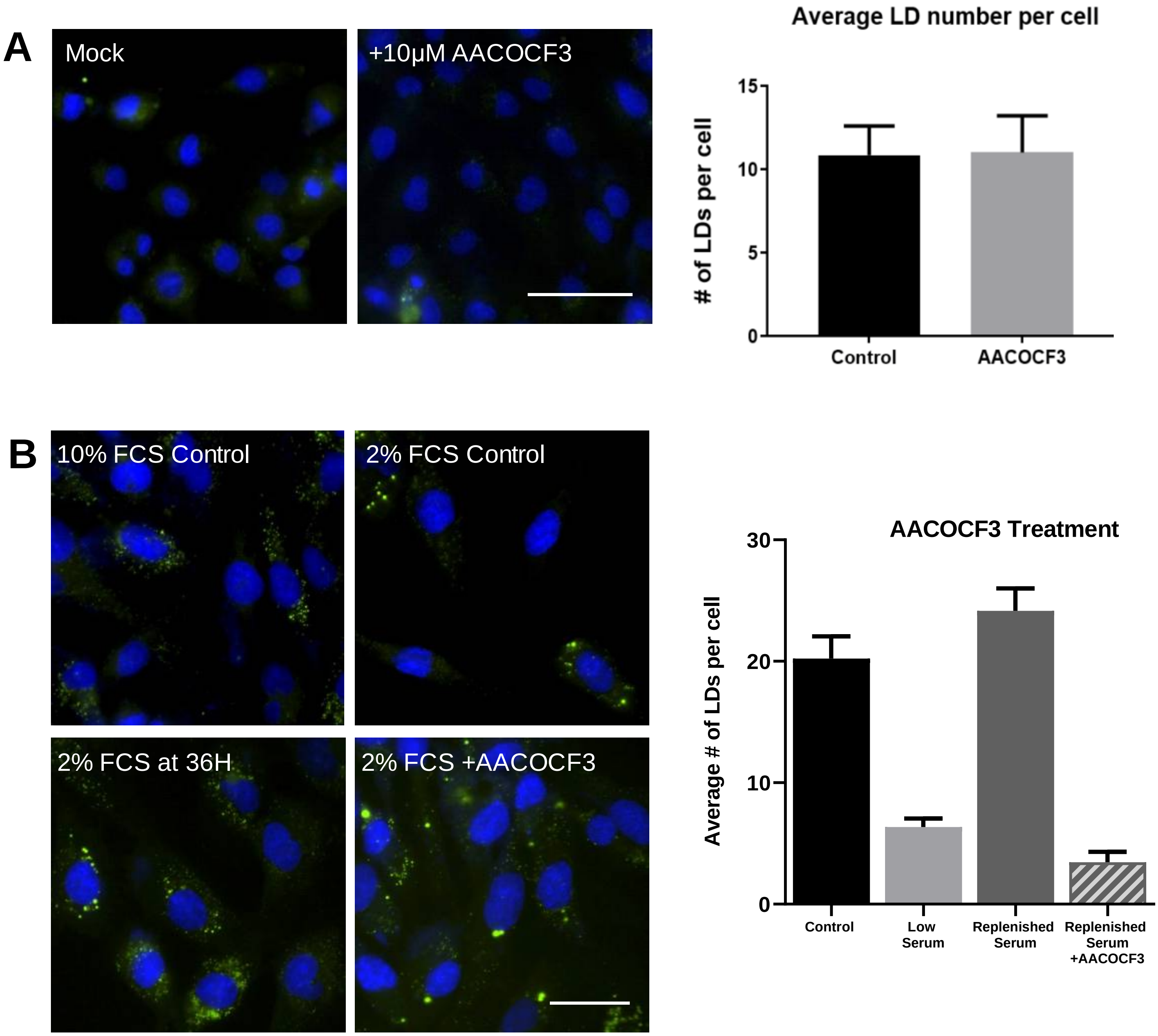

### Supplementary figure 7

#### Supplementary Figure 7

A

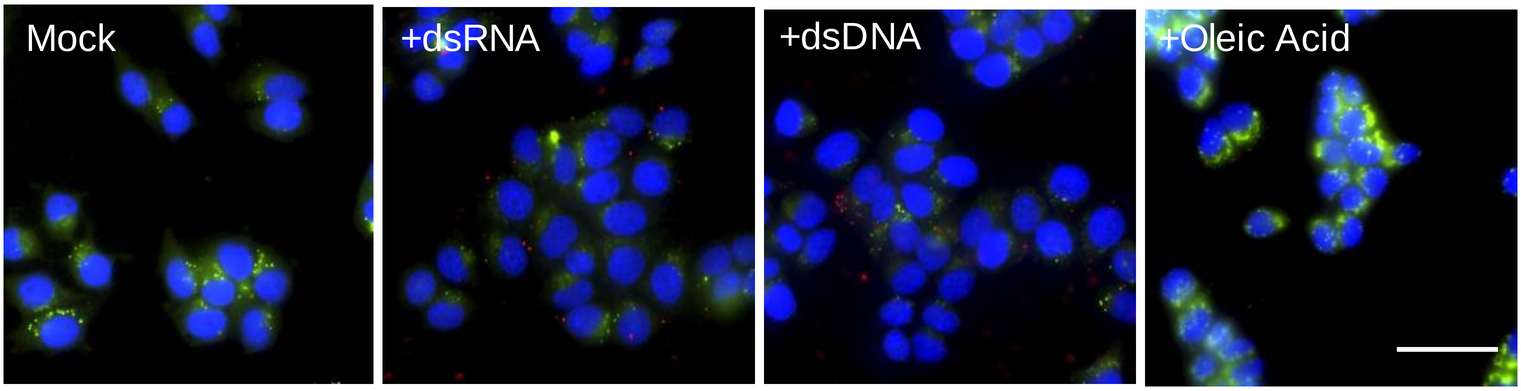


B

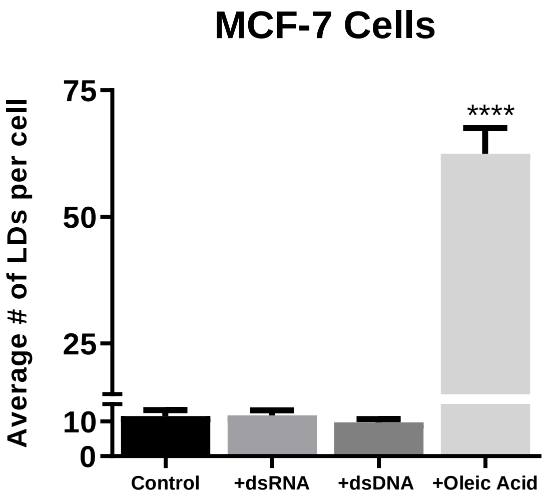


C

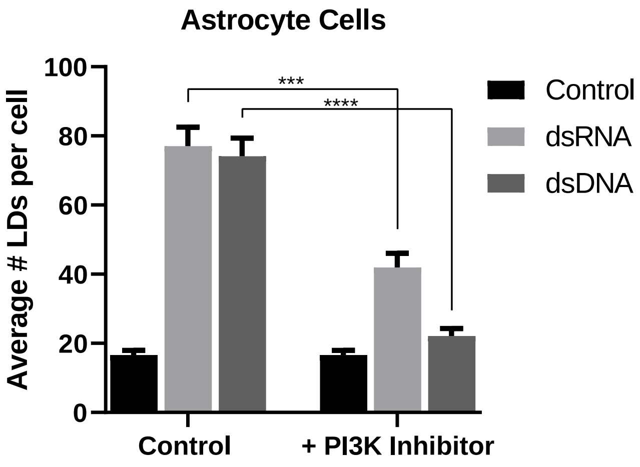

### Supplementary figure 8

#### Supplementary Figure 8

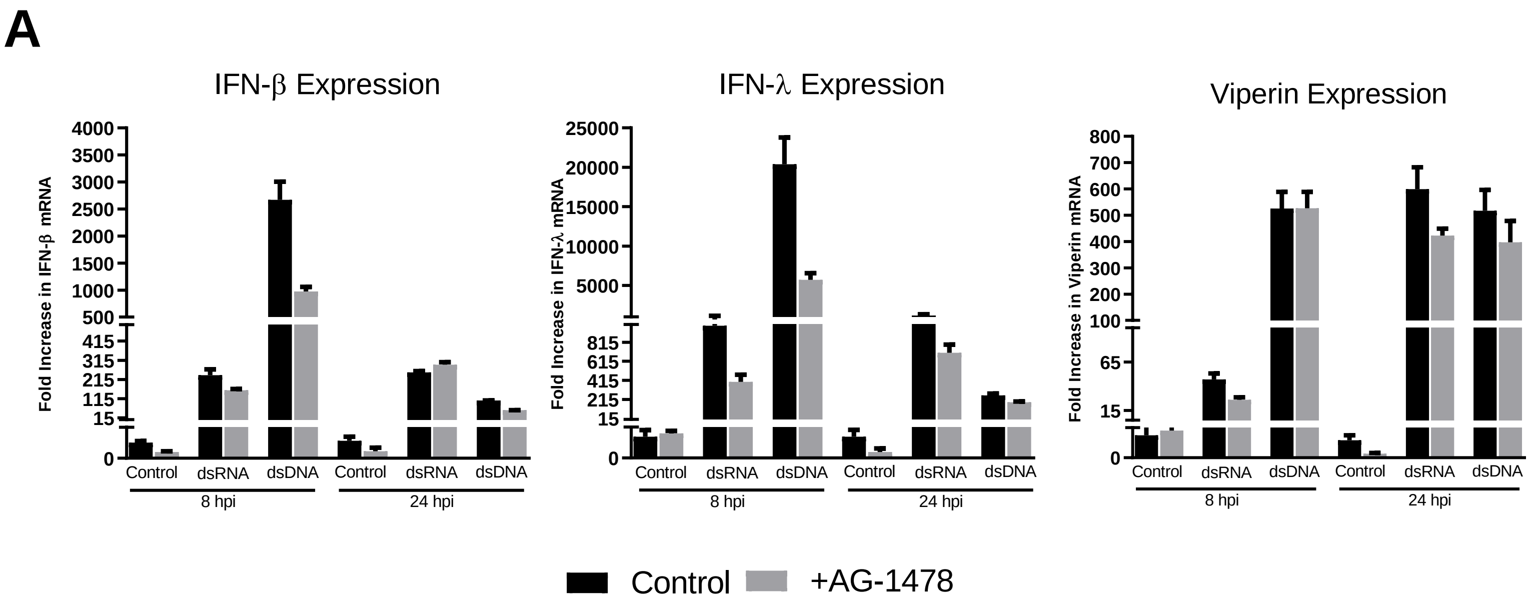

### Supplementary figure 9

#### Supplementary Figure 9

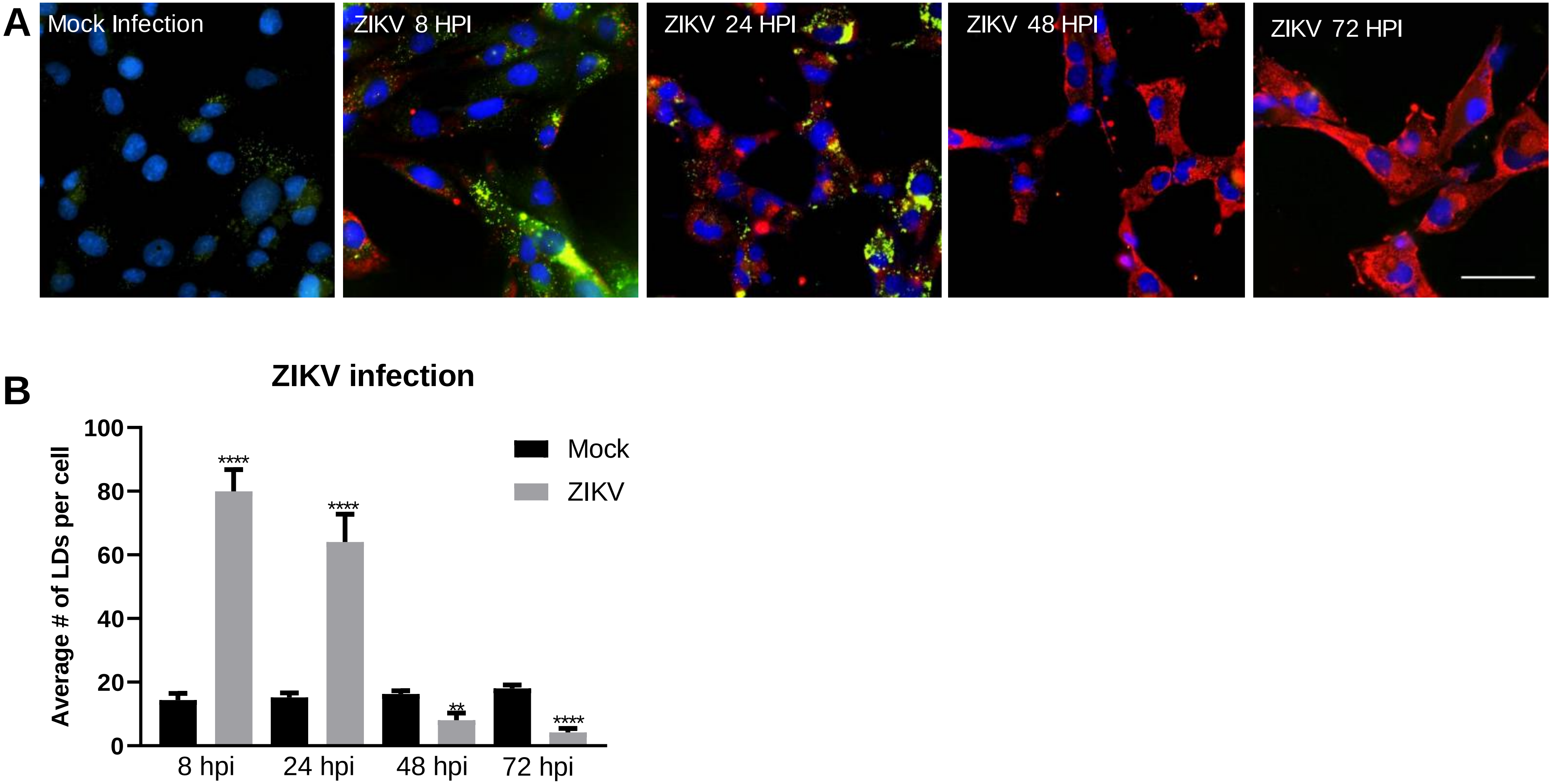
